# Accelerated lung evolution associated with end-Permian and end-Triassic mass extinctions

**DOI:** 10.64898/2026.08.17.745201

**Authors:** Zhili Gu, Zhen Shao, Ziqian Hao, Yi-Hsuan Pan, Haipeng Li

## Abstract

The end-Permian and end-Triassic mass extinctions, driven by massive catastrophic volcanism and prolonged hypoxia, fundamentally reshaped life on Earth. However, the genomic impacts of these two biodiversity crises on living organisms remain largely unknown. Here, we performed a genome-wide screening to identify accelerated evolved regions in the ancestral lineage of mammals that survived both extinction events. Nearly all (20/21) of these accelerated regions were located in protein-coding sequences, and 81% (17/21) were found to be associated with lung function. We further extended our analysis to three additional vertebrate lineages that experienced either one or both of the mass extinctions. Similar genomic signatures, involving accelerated evolution of lung-related genes, were also observed in these non-mammalian lineages. Collectively, these findings suggest that adaptation of lung-related genes to prolonged hypoxia may have occurred during the two mass extinction events, potentially driving a second evolutionary stage of the lung following the vertebrate transition to land.

## Introduction

Mass extinction events have disrupted ecosystem, reduced biodiversity, and promoted taxonomic and morphological diversifications among survived lineages ^1–3^. The end-Permian mass extinction (ePME, also known as Permian–Triassic mass extinction) and the end-Triassic mass extinction (eTME) occurred 252 and 201 million years ago (Mya), respectively. The ePME was triggered by large igneous province (LIP) eruptions of the Siberian Traps, while the eTME resulted from LIP eruptions of the Central Atlantic Magmatic Province ^4–6^. The resulting environmental perturbations, including hypoxia, global warming, aridity, acid rain, wildfires, and ozone destruction driven by these volcanic events, should have been major drivers of terrestrial life loss ^7–9^. The familial extinction rates reached as high as 60.9% and 30.1% for the two events, respectively ^1^, making them two of the “Big Five” mass extinction events ^10^. However, it remains largely unknown how terrestrial animals survived these catastrophic events and how their genomes were affected.

After mammals and birds diverged around 318 Mya, the most recent common ancestor of mammals emerged 180 Mya ^11^. This indicates that the ancestral mammalian lineage was influenced by both mass extinction events (Fig. 1a). The survival of mammalian ancestors through these extinctions may be attributed to their adaptation to drastic environmental changes and the development of mammalian-specific morphological traits ^12^. Under this hypothesis, lineage-specific accelerated evolution in mammals may have been driven by adaptations throughout the Permian, Triassic, and Jurassic periods. Accelerated genes are expected to be enriched in functional categories related to environmental adaptation. Alternatively, the survival of mammalian ancestors could simply be due to refugee habitats, and no specific adaptations to volcanism-related stressors would be expected.

**Fig. 1.**
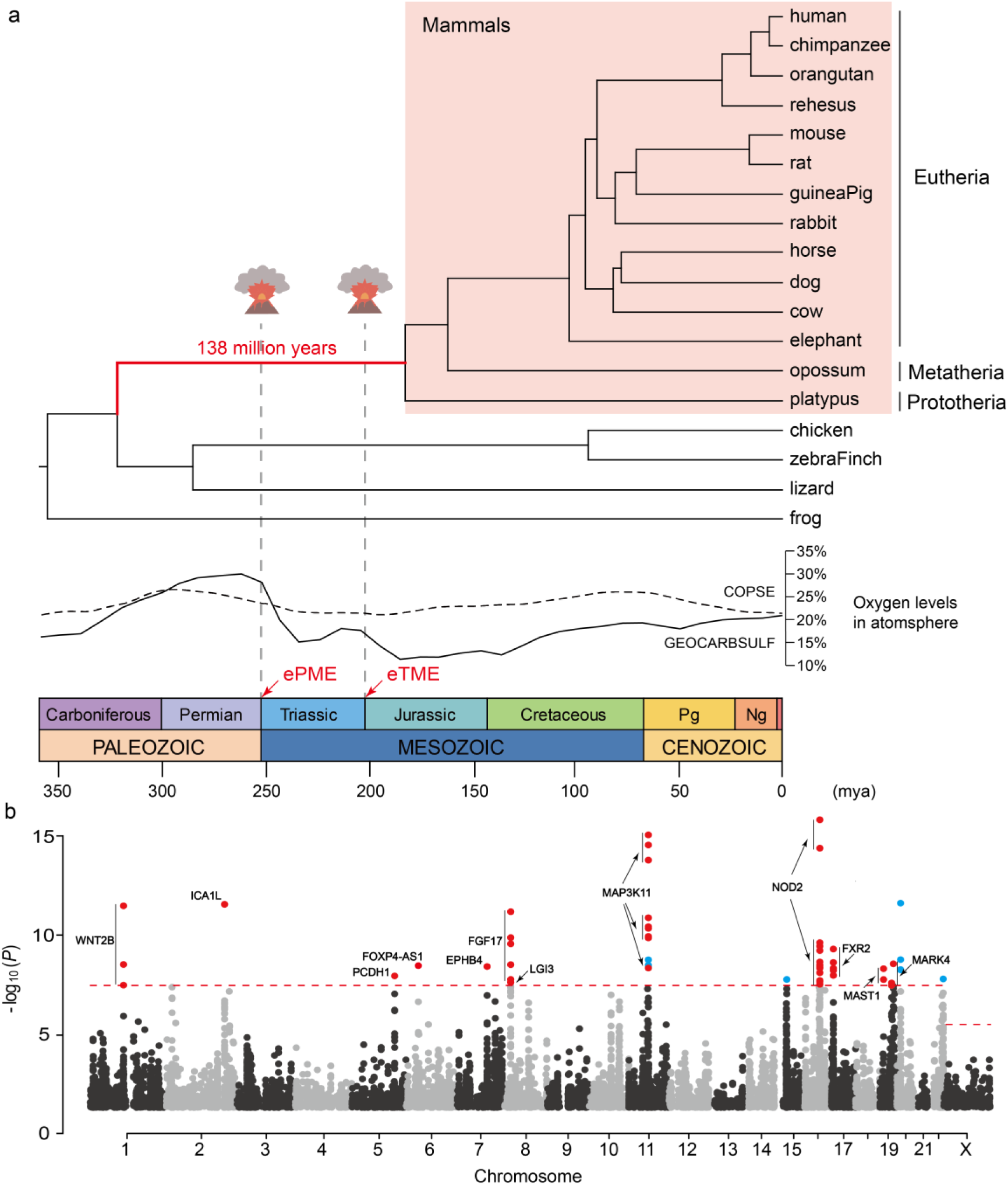
Genome-wide screening for MAWs. **a**, Phylogenetic tree of 18 vertebrates used to detect mammalian accelerated windows (MAWs). The ancestral lineage of mammals is highlighted in red. The end-Permian mass extinction and the end-Triassic mass extinction are indicated above the branch. Estimated atmospheric oxygen concentrations based on the GEOCARBSULF model ^32^ (solid line) and the COPSE model ^33^ (dashed line) are presented below the tree. The chronostratigraphic chart is also shown. ePME: end-Permian mass extinction; eTME: end-Triassic mass extinction; Pg: Paleogene; Ng: Neogene. **b**, Manhattan plot of KFP screening results. A genome-wide scan for sequences with accelerated evolution is presented as a Manhattan plot, showing significance levels against human chromosomal locations. Each dot represents a window with *p* < 0.05. The dash line indicates the significance threshold after Bonferroni correction. Accelerated windows (TOP-MAWs) associated with lung function or lung cancer are highlighted in red, while the others are indicated in blue. Corresponding gene names are displayed adjacent to the dots.

In this study, we screened for accelerated genomic regions in the ancestral mammalian lineage. Out of 1,418,060 windows screened, 11,843 accelerated windows were identified. These accelerated windows showed significant enrichment for genes involved in lung cancers. We also found that nearly all of the top-accelerated regions are located in coding regions, and most are associated with lung function or related diseases. The results suggested an adaptation of the mammalian respiratory system during the ePME and eTME. Moreover, the ancestor of turtles survived both the ePME and eTME, while the ancestor of archosaurs and the ancestor of birds each survived one of these mass extinctions. Accelerated evolution analyses conducted across these three ancestral lineages also revealed lung-related adaptive evolution in all cases. Therefore, our findings implied a link between the volcanism-driven mass extinctions and lung evolution in mammals and other terrestrial vertebrates. These results may also indicate the second lung-related evolutionary stage of terrestrial vertebrate since its water-to-land transition.

## Results

### Genomic screening for mammal-accelerated regions

A total of 18 representative vertebrate species—including 14 mammals, two birds, one reptile, and one amphibian—were selected to identify accelerated evolution specific to ancestral mammals (Fig. 1a). The known species tree and corresponding branch lengths were obtained from TimeTree ^11^. Multi-genome alignments of the 18 species were constructed, and the resulting genome tree was confirmed to be consistent with the known species tree (Fig. S1). Genomic analyses were then performed to detect mammal-accelerated regions using the Kung-Fu Panda (KFP) software ^13^, with a sliding window size of 100 bp and a step size of 20 bp. For each window, evolutionary rates in mammalian and non-mammalian vertebrates were normalized and compared to those of the ancestral mammalian lineage. The traditional approaches ^14–17^ can only detect the accelerated evolution over conserved regions. However, by applying KFP, we were able to detect this over both conserved and non-conserved regions.

In total, 1,418,060 autosomal and 48,911 chr-X windows were analyzed. By comparing the genomic locations of these autosomal windows with the RefSeq human gene annotations ^18^, we found that the analyzed windows are more likely to be located within genes rather than intergenic regions (86.6% *vs* 13.4%). This distribution was expected, as the detection of accelerated evolution in the ancestral mammalian lineage required the presence of homologous fragments across frogs, birds, reptiles, and mammals (Fig. 1a). Consequently, highly variable genomic regions may have been excluded from the analysis due to this requirement.

### Top mammal-accelerated regions enriched in protein coding regions

Of the 1,466,971 windows analyzed, those with a *P*-value < 0.05 showed a weak signal of accelerated evolution in the mammalian ancestor and were therefore designated as mammalian accelerated windows (MAWs). A total of 11,762 MAWs were identified, comprising 11,482 autosomal MAWs and 280 X-chromosomal MAWs. After Bonferroni correction for multiple testing ^19^, 57 autosomal windows remained significant (*P*-value < 3.53 ×10^−8^ and 1.02 ×10^−6^ for autosomes and the X chromosome) (Fig. 1b). These 57 MAWs were labeled as TOP-MAWs, as they represent the most strongly accelerated evolutionary signals in the ancestral mammalian lineage. Furthermore, adjacent MAWs were merged, resulting 5,211 mammalian accelerated regions (MARs), which include 5,064 autosomal MARs and 147 X-chromosomal MARs (Supplemental Excel 1). The average length of MARs is 128 bp. The MARs were ranked according to its smallest *P*-value among corresponding windows. After Bonferroni correction, the 21 most accelerated MARs were designated as TOP-MARs (Table S1).

Based on the genomic locations of the analyzed windows, we found that autosomal MAWs show only a slight bias toward coding sequence (CDS) regions (*P*-value = 0.1408, binomial test), whereas TOP-MAWs are highly significantly enriched in CDS regions (*P*-value = 2.17×10^-12^, binomial test) (Fig. 2a). In particular, 56 out of 57 TOP-MAWs (98.2%) are located within CDS regions, underscoring the importance of adaptive protein evolution.

**Fig. 2.**
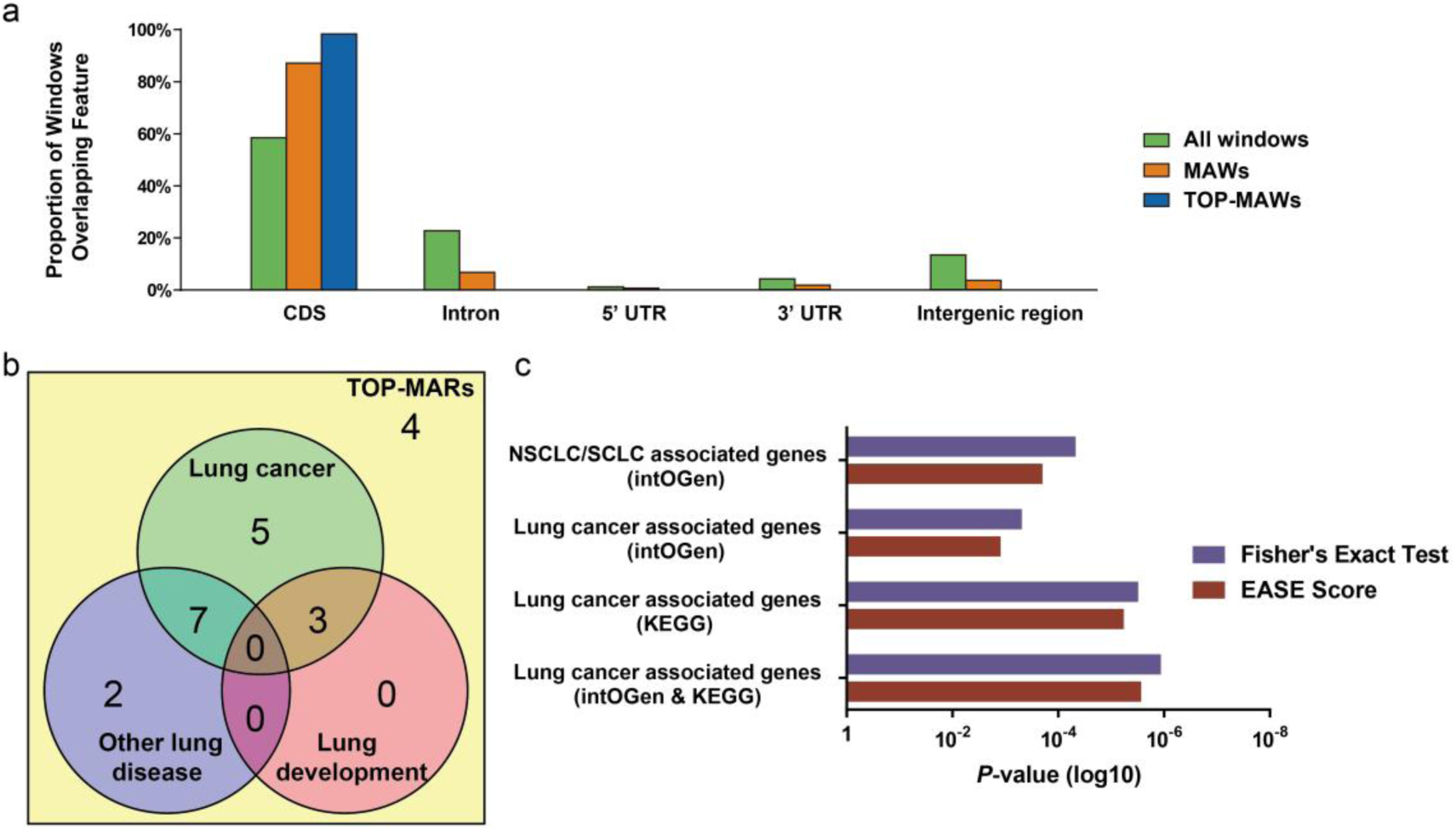
Statistical information of MAWs and TOP-MARs. **a**, Proportion of autosomal windows overlapping genomic features in the human genome. Overlapping feathers are presented for three groups: all windows (1,418,060 windows total), MAWs (11,482 windows total), and TOP-MAWs (57 windows total). **b**, Overlap of TOP-MARs with lung function and diseases. Seventeen out of 21 TOP-MARs are associated with lung function or diseases, including: 15 TOP-MARs associated with lung cancer, 9 with other lung diseases, and 3 with lung development. **c**, Enrichment analysis of MAW-associated genes in lung cancer-related terms. Fisher’s exact test was used to assess gene enrichment in four customized lung cancer-related gene lists from Integrative Onco Genomics (intOGen) and Kyoto Encyclopedia of Genes and Genomes (KEGG) pathways. Results are shown as *P*-values and EASE scores (a variant of the one-tailed Fisher’s exact probability value). NSCLC: non-small cell lung cancer; SCLC: small cell lung cancer.

### Top mammal-accelerated regions related to lung function/diseases

Since 56 of the TOP-MAWs are located within CDS regions, we identified the genes containing these TOP-MAWs and reviewed their functional annotations based on peer-reviewed publications. We found that a very high percentage of TOP-MAWs (51/57, or 89.5%) are associated with genes related to lung function or related diseases (Fig. 1b). To account for potential bias due to gene size, we examined TOP-MARs that were formed by merging neighboring MAWs. Among 21 TOP-MARs, 17 were found to be related to lung function or diseases. The majority (15 out of 17) of these lung-related TOP-MARs are associated with lung cancer (Table S1, Fig. 2b). Furthermore, 21 TOP-MARs correspond to 16 unique genes and one lncRNA, 13 of which are associated with lung function or diseases. For example, the most accelerated gene *NOD2* (nucleotide-binding oligomerization domain 2) is involved in lung inflammation ^20^, *Staphylococcus aureus*-induced pheumonia ^21^ and *Legionella pneumophila* infection ^22^. The gene *MAP3K11*, which harbors TOP-MAR2 and TOP-MAR3, is targeted by microRNAs that promote myofibroblast proliferation in pulmonary fibrosis ^23^ and play an anti-proliferation role in non-small cell lung cancer ^24^. Overall, all these results indicated a lung-related adaptive mammalian evolution.

To further validate the association between mammalian accelerated regions and lung cancer, a total of 3,404 genes related to the 11,482 autosomal MAWs were collected for enrichment analysis. Four categories of lung cancer-related genes (Table S2) were obtained from KEGG (Kyoto Encyclopedia of Genes and Genomes) ^25^ and intOGen (Integrative Onco Genomics) ^26^. The enrichment analyses showed that MAW-associated genes are significantly enriched in all four categories (Fig. 2c). These results confirmed the accelerated evolution of lung function or disease-related genes in the ancestral lineage of mammals, suggesting an adaptation related to volcanism-driven environmental changes.

### Lung-oriented adaptation during ePME and eTME

Our results presented above suggested a volcanism-driven lung-related adaptation in mammalian evolution. We therefore asked whether this can be detected in other terrestrial animal clades because adaptation to volcanism-driven environmental changes should not be limited to the mammalian clade. To investigate this, we examined three non-mammalian ancestral lineages that experienced one or both of the mass extinction events. The ancestral archosaurs (i.e., the common ancestor of birds and crocodylians) survived the ePME (Fig. 3a). The ancestors of birds survived the eTME (Fig. 3b). The ancestors of turtles underwent both the ePME and the eTME (Fig. 3c).

**Fig. 3.**
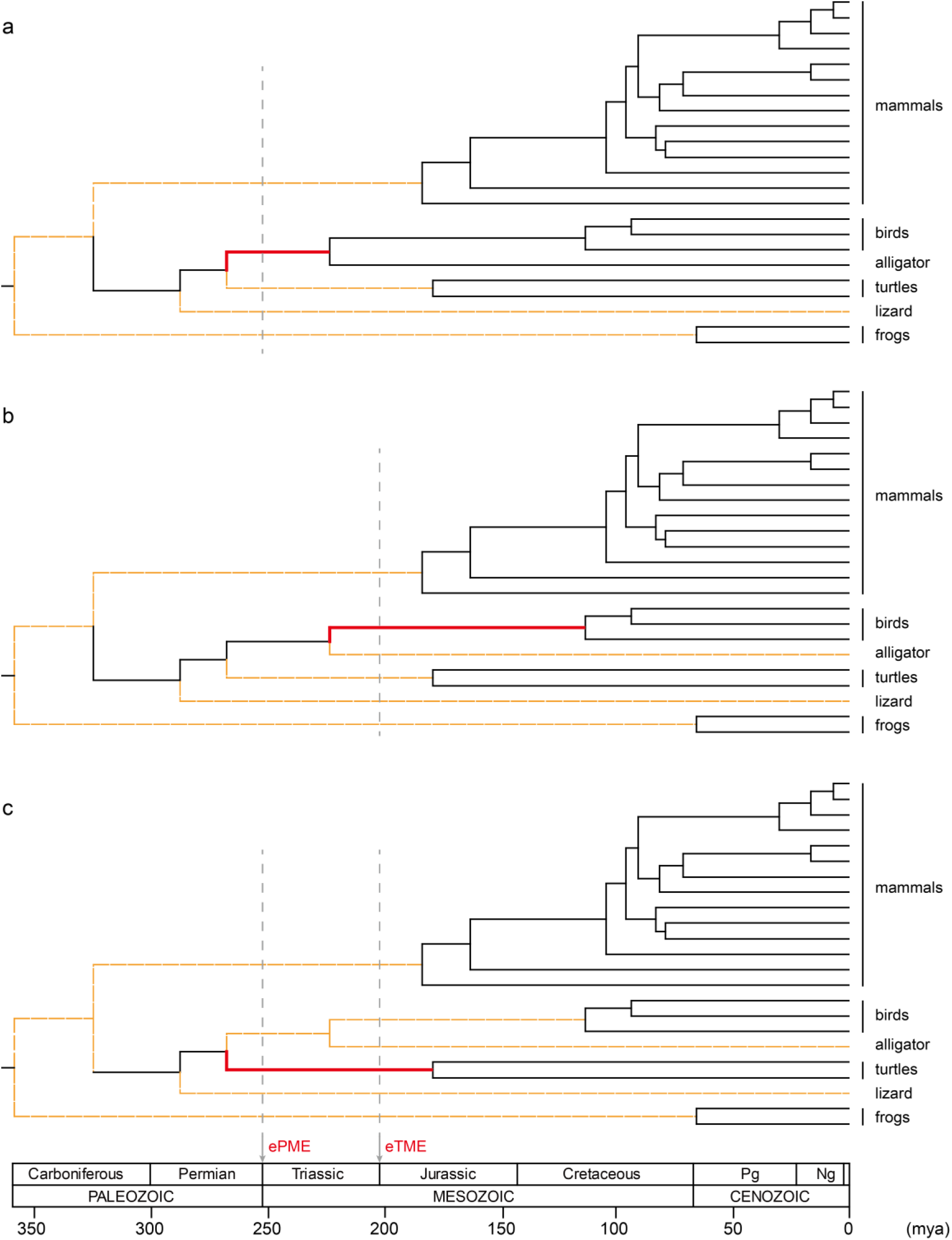
Expanded phylogenetic tree of 23 vertebrates for detecting accelerated regions in the ancestral lineage of archosaurs (**a**), birds (**b**), and turtles (**c**). The red branch indicates the ancestral lineage under analysis. Branches marked with dashed orange lines were excluded when calculating the neutral evolutionary rate. The grey vertical dash line indicates the timing of two mass extinction events, labeled above the global chronostratigraphic chart. ePME: end-Permian mass extinction; eTME: end-Triassic mass extinction; Pg: Paleogene; Ng: Neogene.

After scanning for the accelerated evolution of the three ancestral clades (Fig. 3), the top five accelerated regions and the functions of their corresponding genes were analyzed for each ancestral clade (Table S3). We found that 93% (14/15) of the candidate regions are located in CDS regions, consistent with the results observed in the mammal-specific lineage. Moreover, the top five candidates from each of the three analyses each yielded three genes associated with lung function or diseases. These findings suggested that the accelerated evolution of these lung-related genes is likely due to volcanism-driven adaptation, reflecting a common evolutionary response to extreme environmental conditions following massive volcanic eruptions.

## Discussion

In this study, we discovered that all the four ancestral lineages (Mammlia, Archosauria, Aves, and Testudines) exhibited patterns of accelerated evolution in lung-related genes. Since all the ancestral lineages independently survived the ePME and/or eTME, this strongly suggested that the respiratory system of terrestrial vertebrate was among the most severely affected organs by the adverse environmental conditions following massive volcanic eruptions. Consequently, accelerated evolution in lung-related traits may have provided a selective advantage to these lineages. The proportion of lung-oriented accelerated regions in Archosauria, Aves, and Testudines (60% on average) was less pronounced than that in the mammalian clade (about 80%). We speculate that this difference may be attributed to the higher tolerance to hypoxia in the ancestors of archosaurs, ancestor of birds, and ancestor of turtles ^27,28^.

During massive volcanic eruptions, volcanic ash likely had adverse effects on the ancestors of Mammalia and other taxa. Although these eruptions lasted only shortly, volcanic ash accumulated in the environment and was redistributed across hemispheric scales ^29^. Long-term exposure to volcanic ash can cause acute or chronic respiratory effects such as asthma, pneumoconiosis, and cancer ^30,31^. Therefore, we proposed that selective pressure imposed by volcanic ash may have contributed to accelerated evolution of the respiratory system of terrestrial vertebrate.

After the two massive volcanic eruptions, substantial declines in atmospheric oxygen levels occurred ^32,33^ (Fig. 1a). Hypoxia is therefore widely recognized as a major cause of terrestrial life decline during these volcanic eruption-driven mass extinctions ^34–38^. Hypoxia promotes the production of reactive oxygen species from mitochondria, leading to an imbalance in the redox state in the lung and resulting in conditions such as hypoxic vasoconstriction, pulmonary edema, and chronic pulmonary hypertension ^39^. Prolonged low oxygen levels would have been lethal to many species, either driving them to extinction or forcing them to adapt the harsh conditions. Therefore, adaptive evolution of the lung was likely driven by long-term hypoxia resulting from the two massive volcanic eruptions (*i.e.*, the ePME and eTME).

The vertebrate water-to-land transition (419–359 Mya) was a critical period for adaptive lung evolution ^40,41^. Interestingly, the lung-related accelerated regions identified in this study are not linked to known genetic innovations associated with lung adaptation in early ray-finned fishes ^42^ or the African lungfish ^43^. Instead of being associated with broad pulmonary phenotypes, these accelerated regions show significant enrichment in lung cancer and other lung diseases. Based on these findings, we propose a two-stage model of lung evolution. The first stage involved adaptation of the respiratory apparatus in the ancestors of tetrapods around 419–359 Mya, driven by atmospheric oxygenation changes that facilitated the water-to-land transition ^44^. The second stage consisted of a subtle, lung-oriented accelerated evolution that occurred independently in various terrestrial lineages, such as mammalian and avian ancestors, between about 260 and 190 Mya. The later adaptation was probably a response to prolonged low oxygen conditions, which drove the adaptive changes of lung-related genes.

The volcanism-related adaptation in the different ancestral lineages of terrestrial animals may not be exclusively due to the two massive volcanic eruptions. In addition to the devastating ePME and eTME (Fig. 1a), two other lesser volcanism-related extinctions (i.e., Smithian/Spathian boundary event and Carnian pluvial event) occurred around 249 and 234 Mya ^45,46^ (Table S4). All four known biodiversity crises have been linked to massive volcanic eruptions, which led to similarly severe environmental impacts ^47^. Overall, these major and lesser extinction events may have provided prolonged and recurrent global selective pressures that shaped the lung evolution in the ancestral lineage of mammals.

In conclusion, this study highlights that accelerated evolution of lung-related genes may have enabled ancestral lineages, especially ancestral mammals, to survive multiple volcanism-related mass extinctions and adapt to prolonged hypoxia during the Triassic and Jurassic periods. This accelerated evolution is enriched in genes associated with lung cancer. Therefore, our findings suggest that volcanism-driven mass extinctions not only played a significant role in the evolution of terrestrial animals, but also may contribute to the underlying causes of lung cancer in humans.

## Materials and Methods

### Multi-genome alignments

In this study, we chose 18 representative vertebrate species when detecting the accelerated evolution on the ancestral lineage of mammals. The species set includes 12 eutherians, one metatherian, one prototherian, two birds, one reptile, and one amphibian. Those 18 species are: *Homo sapiens* (human, GRCh38/hg38), *Pan troglodytes* (chimpanzee, CSAC 2.1.4/panTro4), *Pongo abelii* (orangutan, WUGSC 2.0.2/ponAbe2), *Macaca mulatta* (rhesus, CR_1.0/rheMac3), *Mus musculus* (mouse, GRCm38/mm10), *Rattus norvegicus* (rat, RGSC 6.0/rn6), *Cavia porcellus* (guinea pig, Broad/cavPor3), *Oryctolagus cuniculus* (rabbit, Broad/oryCun2), *Bos taurus* (cow, UMD_3.1.1/bosTau8), *Equus caballus* (horse, Broad/equCab2), *Canis lupus familiaris* (dog, Broad CanFam3.1/canFam3), *Loxodonta africana* (elephant, Broad/loxAfr3), *Monodelphis domestica* (opossum, Broad/monDom5), *Ornithorhynchus anatinus* (platypus, WUGSC 5.0.1/ornAna1), *Gallus gallus* (chicken, ICGSC Gallus_gallus-4.0/galGal4), *Taeniopygia guttata* (zebra finch, WashU taeGut324/taeGut2), *Anolis carolinensis* (lizard, Broad AnoCar2.0/anoCar2), and *Xenopus tropicalis* (western clawed frog, JGI 7.0/xenTro7).

Genome sequences of those 18 species and their pairwise genome alignments with the human genome as the reference were downloaded from the UCSC genome browser website (http://hgdownload.soe.ucsc.edu/downloads.html). UCSC Kent utilities (http://hgdownload.cse.ucsc.edu/admin/jksrc.zip) were used for fetching or generating information of chromosome size. Multi-genome alignments of 18 species were created by MULTIZ ^48^ (http://www.bx.psu.edu/miller_lab/).

To detect the accelerated evolution on the ancestral lineages of turtles, archosaurs and birds, the species set was expanded to include five additional species. The five additional species are: *Apteryx mantelli* (brown kiwi, MPI-EVA AptMant0/aptMan1), *Alligator mississippiensis* (American alligator, allMis0.2/allMis1), *Chelonia mydas* (green seaturtle, CheMyd_1.0/cheMyd1), *Pelodiscus sinensis* (Chinese softshell turtle, PelSin_1.0/pelSin1), and *Xenopus laevis* (African clawed frog, Xenopus_laevis_v2/xenLae2). Multi-genome alignments of expanded species set (23 species) were obtained using the methods described above.

### Phylogenetic tree

The phylogenetic trees were obtained by using the Common Tree function of the NCBI Taxonomy database (https://www.ncbi.nlm.nih.gov/Taxonomy/CommonTree/wwwcmt.cgi). Median divergence time of each species pair was obtained from the TimeTree website (http://www.timetree.org) ^11^ and used as the branch length of the phylogenetic trees. The phylogenetic tree of the 18 species (Fig. 1a) was provided in Newick format as follows:

((((((((((human:6.4,chimpanzee:6.4):8.8,orangutan:15.2):13.61,rhesus:28.81): 60.19,(((mouse:15.9,rat:15.9):54.1,guineaPig:70):10,rabbit:80):9):5,(cow:81,(horse:7 7.1,dog:77.1):3.9):13):8,elephant:102):58,opossum:160):20,platypus:180):138,((chic ken:92.9,zebraFinch:92.9):189.1,lizard:282):36):33.7,frog:351.7):0.

The phylogenetic tree of the expanded species set (Fig. 3) is ((((((((((human:6.4,chimpanzee:6.4):8.8,orangutan:15.2):13.61,rhesus:28.81): 60.19,(((mouse:15.9,rat:15.9):54.1,guineaPig:70):10,rabbit:80):9):5,(cow:81,(horse:7 7.1,dog:77.1):3.9):13):8,elephant:102):58,opossum:160):20,platypus:180):138,(((((c hicken:92.9,zebraFinch:92.9):20,brownKiwi:111.2):108,alligator:219.2):42.8,(green Seaturtle:175,chineseSoftshellturtle:175):87):20,lizard:282):36):33.7,(westernClawe dfrog:64,africanClawedfrog:64):287.7):0.

### Identification of accelerated genomic regions

With the multi-genome alignments and phylogenetic trees, we used the Kung-Fu Panda (KFP) software ^13^ to scan for accelerated sequences in the ancestral lineage of mammals (*i.e.*, the red branch in Fig. 1a). A sliding window analysis was conducted with a window size of 100 bp and a step size of 20 bp. For each window, the following criteria were applied. (a) At least half of the species had to contain normal sites (excluding indels and low-quality sequences). (b) The number of missing species had to be no more than four. (c) The percentage of indels among non-missing species should be less than 30%. Windows with a *P*-value below 0.05 were identified as mammalian accelerated windows (MAWs) before correcting for multiple tests. Adjacent accelerated windows were then merged, and the resulting mammalian accelerated regions (MARs) were ranked according to the lowest *P*-value within each merged region. To obtain statistically significant accelerated windows, a Bonferroni correction ^19^ was applied to control the family-wise error rate at 0.05. The adjusted significance thresholds were set at 3.53 ×10^−8^ for autosomal and 1.02 ×10^−6^ for X-chromosomal TOP-MAWs and TOP-MARs.

The KFP software was also used to detect accelerated evolution in the ancestral lineages of archosaurs, birds, and turtles (*i.e.*, the red branches in Fig. 3). Since multiple lineages were affected by one or two major extinction events, we accounted for this effect in our subsequent analysis. For example, the ancestral lineage of archosaurs was impacted by the ePME. Therefore, when detecting accelerated evolution in this lineage, four internal/external lineages (indicated by dashed orange lines in Fig. 3) were excluded from the analysis, as they were also influenced by the ePME. This approach ensured that the neutral evolutionary rate of each examined fragment was estimated based on the remaining lineages (marked by solid black lines in Fig. 3).

### Enrichment analysis

To perform the enrichment analysis, we adopted a widely used method to generate the reference gene list by considering all analyzed windows. If a window overlapped with a gene, the corresponding gene name was recorded. For windows located in intergenic regions, flanking genes were identified using the Galaxy platform ^49,50^. These genes were compiled into the reference gene list to correct for sampling bias in the enrichment analysis. Using this approach, we also derived the list of genes associated with mammalian accelerated windows (MAWs).

We then constructed four optimized categories for lung-related genes by merging lung-related gene sets from KEGG (Kyoto Encyclopedia of Genes and Genomes) ^25^ and intOGen (Integrative Onco Genomics) ^26^. For NSLC/SCLC associated genes (intOGen), we included mutational cancer driver genes of the two most common forms of lung cancer (i.e., non-small cell lung cancer and small cell lung cancer) from the intOGen database. For lung cancer-associated genes (intOGen), we incorporated mutational cancer driver genes from multiple types of lung cancer (including non-small cell lung cancer, small cell lung cancer, lung adenocarcinoma, lung squamous cell carcinoma, and lung neuroendocrine cancer) from the intOGen database. For lung cancer-associated genes (KEEG), we combined genes from the non-small cell lung cancer pathway and the small cell lung cancer pathway from the KEGG pathway database. For lung cancer-associated genes (intOGen & KEEG), we merged the three gene categories described above. The merged gene lists were filtered using the reference gene list and served as target categories for the gene enrichment test.

Fisher’s Exact test and the EASE score (a variant of the one-tailed Fisher’s exact probability value) ^51^ were used to measure gene enrichment in annotation terms.

## Acknowledgement

We would like to thank Profs. Ya-Ping Zhang and Shu-Yi Zhang for their instructive discussions and inspiration 30 years ago on the evolution of mammals.

