## Supplementary material for "Accelerated lung evolution associated with end-Permian and end-Triassic mass extinctions": Fig. S1, Tables S1-S4

**Short title:** Accelerated evolution associated with mass extinctions

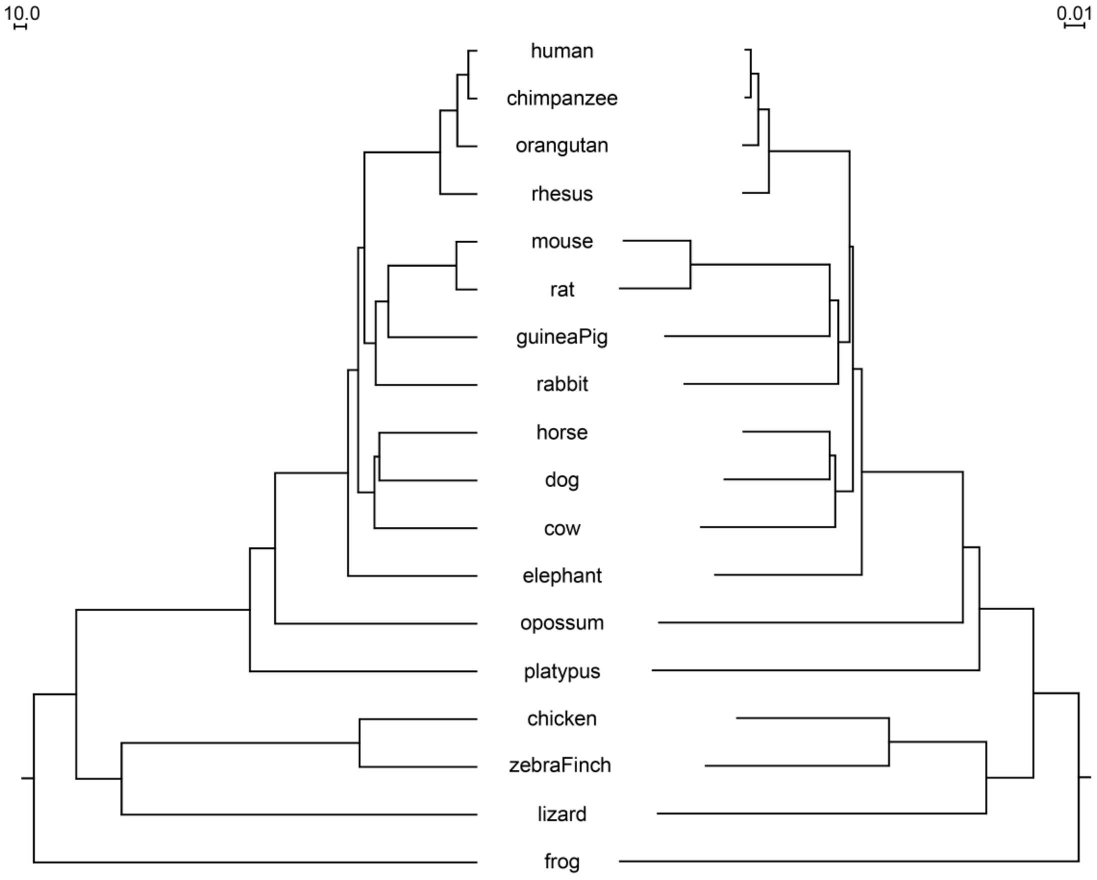


**Fig. S1** Comparison between the original phylogenetic tree (left) and the genome-wide neighbor-joining tree (right) [1] generated by the eGPS software [2].

**Table S1.** List of the accelerated regions after Bonferroni correction in the ancestral lineage of mammals.

| Rank | Name | Coordinate (hg38) | Genomic Location | Associated Gene | *P*-value | Lung related function/disease | Reference |
|---|---|---|---|---|---|---|---|
| 1 | TOP-MAR1 | chr16:50,710,721-50,711,520 | CDS | *NOD2* [a] | $1.54\times10^{-16}$ | lung inflammation, pneumonia | 3-5 |
| 2 | TOP-MAR2 | chr11:65,606,661-65,606,860 | CDS | *MAP3K11* | $8.99\times10^{-16}$ | NSCLC, pulmonary fibrosis | 6,7 |
| 3 | TOP-MAR3 | chr11:65,607,601-65,607,840 | CDS | *MAP3K11* | $1.68\times10^{-14}$ | NSCLC, pulmonary fibrosis | 6,7 |
| 4 | TOP-MAR4 | chr20:13,865,301-13,865,560 | CDS | *SEL1L2* | $2.48\times10^{-12}$ | | |
| 5 | TOP-MAR5 | chr2:202,821,301-202,821,520 | CDS | *ICA1L* | $2.83\times10^{-12}$ | NSCLC | 8 |
| 6 | TOP-MAR6 | chr1:112,517,101-112,517,420 | CDS | *WNT2B* | $3.36\times10^{-12}$ | NSCLC, CPAM | 9,10 |
| 7 | TOP-MAR7 | chr8:22,047,941-22,048,260 | CDS | *FGF17* | $6.65\times10^{-12}$ | pulmonary trunk development, SCLC | 11,12 |
| 8 | TOP-MAR8 | chr11:65,634,101-65,634,380 | CDS | *PCNX3* | $4.46\times10^{-11}$ | | |
| 9 | TOP-MAR9 | chr17:7,592,701-7,592,920 | CDS | *FXR2* | $5.11\times10^{-10}$ | pulmonary hypertension, lung cancer | 13,14 |
| 10 | TOP-MAR10 | chr16:50,712,001-50,712,440 | CDS | *NOD2* | $6.06\times10^{-10}$ | lung inflammation, pneumonia | 3-5 |
| 11 | TOP-MAR11 | chr19:45,302,381-45,302,720 | CDS | *MARK4* | $2.83\times10^{-09}$ | lung cancer, acute lung injury | 15,16 |
| 12 | TOP-MAR12 | chr8:22,046,041-22,046,340 | CDS | *FGF17* | $3.14\times10^{-09}$ | pulmonary trunk development, SCLC | 11,12 |
| 13 | TOP-MAR13 | chr6:41,524,081-41,524,260 | lncRNA | *FOXP4-AS1* | $3.48\times10^{-09}$ | NSCLC | 17 |

| | | | | | | | |
|---|---|---|---|---|---|---|---|
| 14 | TOP-MAR14 | chr7:100,805,141-100,805,320 | CDS | *EPHB4* | $3.89\times10^{-09}$ | NSCLC, lung development | 18-20 |
| 15 | TOP-MAR15 | chr19:12,867,741-12,867,920 | CDS | *MAST1* | $4.99\times10^{-09}$ | LUAD | 21 |
| 16 | TOP-MAR16 | chr17:7,605,621-7,605,800 | CDS | *FXR2* | $1.09\times10^{-08}$ | pulmonary hypertension, lung cancer | 13,14 |
| 17 | TOP-MAR17 | chr5:141,854,161-141,854,360 | CDS | *PCDH1* | $1.16\times10^{-08}$ | NSCLC, Asthma | 22,23 |
| 18 | TOP-MAR18 | chr22:50,177,201-50,178,400 | CDS | *PANX2* | $1.65\times10^{-08}$ | | |
| 19 | TOP-MAR19 | chr15:41,885,761-41,886,400 | CDS | *SPTBN5* | $1.72\times10^{-08}$ | | |
| 20 | TOP-MAR20 | chr8:22,151,441-22,151,680 | CDS | *LGI3* | $2.40\times10^{-08}$ | NSCLC | 24 |
| 21 | TOP-MAR21 | chr19:39,826,641-39,827,000 | CDS | *DYRK1B* | $2.59\times10^{-08}$ | NSCLC | 25,26 |

NSCLC: non-small cell lung cancer; CPAM: congenital pulmonary airway malformations; SCLC: small cell lung cancer; LUAD: lung adenocarcinoma.

[a] Also known as *NLRC2* (NOD-leucine-rich repeat family with caspase recruitment domain 2).

[b] TOP-MAR13 is located in a lncRNA *FOXP4-AS1*. *FOXP4* is the nearest coding gene, also associated with NSCLC.

**Table S2.** Four optimized lists of lung cancer-related genes derived from KEGG (Kyoto Encyclopedia of Genes and Genomes) & intOGen (Integrative Onco Genomics).

| | |
|---|---|
| NSCLC/SCLC associated genes (intOGen) | TP53 LRP1B EGFR KRAS FAT4 KEAP1 PTPRD ARID1A STK11 KMT2D CDKN2A RB1 SMARCA4 ALK ERBB2 PIK3CA ATM RBM10 APC BRAF NF1 ARID2 CUX1 NOTCH1 SETD2 PTPRK CTNNB1 PTEN CMTR2 CUL3 PRF1 ARHGEF10 MAP2K1 FH U2AF1 IDH1 B2M KLF4 BAP1 CTCF CREBBP RUNX1T1 KMT2C EP300 KDM6A LIFR BCR NCOR1 EML4 SET |
| Lung cancer associated genes (intOGen) | TP53 LRP1B EGFR KRAS FAT4 KEAP1 PTPRD ARID1A STK11 KMT2D CDKN2A RB1 SMARCA4 ALK ERBB2 PIK3CA ATM RBM10 APC BRAF NF1 ARID2 CUX1 NOTCH1 SETD2 PTPRK CTNNB1 PTEN CMTR2 CUL3 PRF1 ARHGEF10 MAP2K1 FH U2AF1 IDH1 B2M KLF4 BAP1 CTCF CREBBP RUNX1T1 KMT2C EP300 KDM6A LIFR BCR NCOR1 EML4 SET MGA KDR CDH10 RET RGS7 FLT4 BTK EPHA7 ATF7IP MB21D2 NCOA2 CIC CDC73 EZH2 LRIG3 CLIP1 NRAS PBRM1 SF3B1 MEN1 NFE2L2 FAT1 FN1 RASA1 ARHGAP35 NSD1 FBXW7 DROSHA PRKCB MAML2 FAM135B FGFR3 FGFR2 RAD21 KLF5 HRAS POLQ SUSD2 USP6 CDH11 PABPC1 ARID1B ZNF521 NRK EBF1 TNC LATS2 HLA-A FAT3 RHPN2 NPEPPS SALL4 STAG2 PDGFRA USP8 ESR1 RNF213 |
| Lung cancer associated genes (KEGG) | FHIT RARB RXRA RXRB RXRG CDKN2A CDK4 CDK6 CCND1 RB1 E2F1 E2F2 E2F3 KRAS RASSF1 RASSF5 STK4 PIK3CA PIK3CD PIK3CB PIK3R1 PIK3R2 PIK3R3 PDPK1 AKT1 AKT2 AKT3 BAD CASP9 FOXO3 EGF TGFA EGFR ERBB2 HGF MET GRB2 SOS1 SOS2 HRAS NRAS ARAF BRAF RAF1 MAP2K1 MAP2K2 MAPK1 MAPK3 PLCG1 PLCG2 PRKCA PRKCB PRKCG TP53 CDKN1A GADD45A GADD45B GADD45G BAX BAK1 DDB2 POLK EML4 ALK JAK3 STAT3 STAT5A STAT5B BCL2 CYCS APAF1 CASP3 MYC MAX ZBTB17 CDKN2B CKS1B CKS2 SKP2 CDKN1B CDK2 CCNE1 CCNE2 COL4A2 COL4A4 COL4A6 COL4A1 COL4A5 COL4A3 LAMA1 LAMA2 LAMA3 LAMA5 LAMA4 LAMB1 LAMB2 LAMB3 LAMB4 LAMC1 LAMC2 LAMC3 FN1 ITGA2 ITGA2B ITGA3 ITGA6 ITGAV ITGB1 PTK2 PTEN CHUK IKBKB IKBKG NFKBIA NFKB1 RELA BIRC2 BIRC3 XIAP BIRC7 BCL2L1 TRAF1 TRAF2 TRAF3 TRAF4 TRAF5 TRAF6 PTGS2 NOS2 |
| Lung cancer associated genes (intOGen & KEGG) | TP53 LRP1B EGFR KRAS FAT4 KEAP1 PTPRD ARID1A STK11 KMT2D CDKN2A RB1 SMARCA4 ALK ERBB2 PIK3CA ATM RBM10 APC BRAF NF1 ARID2 CUX1 NOTCH1 SETD2 PTPRK CTNNB1 PTEN CMTR2 CUL3 PRF1 ARHGEF10 MAP2K1 FH U2AF1 IDH1 B2M KLF4 BAP1 CTCF CREBBP RUNX1T1 KMT2C EP300 KDM6A LIFR BCR NCOR1 EML4 SET MGA KDR CDH10 RET RGS7 FLT4 BTK EPHA7 ATF7IP MB21D2 NCOA2 CIC CDC73 EZH2 LRIG3 CLIP1 NRAS PBRM1 SF3B1 MEN1 NFE2L2 FAT1 FN1 RASA1 ARHGAP35 NSD1 FBXW7 DROSHA PRKCB MAML2 FAM135B FGFR3 FGFR2 RAD21 KLF5 HRAS POLQ SUSD2 USP6 CDH11 PABPC1 ARID1B ZNF521 NRK EBF1 TNC LATS2 HLA-A FAT3 RHPN2 NPEPPS SALL4 STAG2 PDGFRA USP8 ESR1 RNF213 FHIT RARB RXRA RXRB RXRG CDK4 CDK6 CCND1 E2F1 E2F2 E2F3 RASSF1 RASSF5 STK4 PIK3CD PIK3CB PIK3R1 PIK3R2 PIK3R3 PDPK1 AKT1 AKT2 AKT3 BAD CASP9 FOXO3 EGF TGFA HGF MET GRB2 |

| | |
|---|---|
| | SOS1 SOS2 ARAF RAF1 MAP2K2 MAPK1 MAPK3 PLCG1 PLCG2 PRKCA PRKCG CDKN1A GADD45A GADD45B GADD45G BAX BAK1 DDB2 POLK JAK3 STAT3 STAT5A STAT5B BCL2 CYCS APAF1 CASP3 MYC MAX ZBTB17 CDKN2B CKS1B CKS2 SKP2 CDKN1B CDK2 CCNE1 CCNE2 COL4A2 COL4A4 COL4A6 COL4A1 COL4A5 COL4A3 LAMA1 LAMA2 LAMA3 LAMA5 LAMA4 LAMB1 LAMB2 LAMB3 LAMB4 LAMC1 LAMC2 LAMC3 ITGA2 ITGA2B ITGA3 ITGA6 ITGAV ITGB1 PTK2 CHUK IKBKB IKBKG NFKBIA NFKB1 RELA BIRC2 BIRC3 XIAP BIRC7 BCL2L1 TRAF1 TRAF2 TRAF3 TRAF4 TRAF5 TRAF6 PTGS2 NOS2 |

**Table S3.** Top five accelerated evolved regions and their associated genes in the ancestral lineage of archosaurs, birds, and turtles.

| **Accelerated evolved branch** | **Massive extinction involved** | **Rank** | **Coordinates** | ***P*-value** | **Related Gene** | **Genomic Location** | **Lung related function / disease** | **Reference** | **Ratio of Lung related regions** |
|---|---|---|---|---|---|---|---|---|---|
| Archosaurs | ePME | 1 | chr11:3,634,241-3,634,500 | $6.49\times10^{-33}$ | TRPC2 | CDS | - | | 3/5 |
| | | 2 | chr11:66,471,181-66,471,420 | $1.84\times10^{-24}$ | PELI3 | CDS | NSCLC | 27,28 | |
| | | 3 | chr5:35,965,341-35,965,920 | $6.63\times10^{-24}$ | UGT3A1 | CDS | - | | |
| | | 4 | chr14:21,393,041-21,393,300 | $1.19\times10^{-23}$ | CHD8 | CDS | LUAD | 29 | |
| | | 5 | chr12:40,433,721-40,433,940 | $3.85\times10^{-23}$ | MUC19 | CDS | HMPV infection | 30 | |
| Birds | eTME | 1 | chr3:71,244,581-71,244,940 | $2.94\times10^{-28}$ | FOXP1 | intron | LUAD, lung development | 31,32 | 3/5 |
| | | 2 | chr19:13,090,281-13,090,400 | $2.44\times10^{-25}$ | NFIX | CDS | acute lung injury, lung cancer | 33-35 | |
| | | 3 | chr2:70,704,261-70,704,480 | $1.47\times10^{-24}$ | ADD2 | CDS | - | | |
| | | 4 | chr11:62,630,361-62,630,520 | $5.17\times10^{-24}$ | GABAB | CDS | - | | |
| | | 5 | chr1:225,144,361-225,144,580 | $1.27\times10^{-23}$ | DNAH14 | CDS | obstructive lung disease | 36 | |
| Turtles | ePME eTME | 1 | chr14:53,950,461-53,950,920 | $1.39\times10^{-31}$ | BMP4 | CDS | pulmonary fibrosis, LUAD, lung development | 37-39 | 3/5 |
| | | 2 | chr20:46,174,501-46,174,720 | $3.79\times10^{-29}$ | CDH22 | CDS | - | | |
| | | 3 | chr4:143,185,421-143,185,780 | $2.90\times10^{-24}$ | USP38 | CDS | | | |
| | | 4 | chr12:57,173,761-57,173,980 | $7.32\times10^{-21}$ | LRP1 | CDS | lung inflammation, | 40,41 | |

| | | | | | | |
|---|---|---|---|---|---|---|
| | | | | | LUAD | |
| 5 | chr6:30,740,641-30,740,840 | $9.24 \times 10^{-21}$ | FLOT1 | CDS | LUAD, SCLC, NSCLC | 42-44 |

NSCLC: non-small cell lung cancer; HMPV: human metapneumovirus; LUAD: lung adenocarcinoma; SCLC: small cell lung cancer.

**Table S4.** Detailed information of mass extinction events during the evolution of the mammalian ancestor (partially adapted from [45]).

| **Extinction** | **Geological time** | **Associated LIP** | **Associated impact structure** | **Global warming or cooling?** | **$\delta^{13}C$** | **Note** |
|---|---|---|---|---|---|---|
| end-Permian | Changhsingian (Lopingian, Permian) | Siberian Traps | None | Yes (+10 ℃) | up to −8‰ | "Big five" |
| Smithian/Spathian | Olenekian (Lower Triassic) | Siberian Traps | None | Yes (+6 ℃) | −6‰ followed by +6‰ | |
| Carnian | Carnian (Upper Triassic) | Wrangellia | None | Yes (+7 ℃) | −5‰ | |
| end-Triassic | Rhaetian (Upper Triassic) | CAMP | None | Yes (+6 ℃) | −5‰ | "Big five" |

CAMP: central Atlantic magmatic province.
